# Menopause status and within group differences in chronological age affect the functional neural correlates of spatial context memory in middle-aged females

**DOI:** 10.1101/2023.09.06.556387

**Authors:** Arielle Crestol, Sricharana Rajagopal, Rikki Lissaman, Annalise LaPlume, Stamatoula Pasvanis, Rosanna K. Olsen, Gillian Einstein, Emily Jacobs, M. Natasha Rajah

## Abstract

Reductions in the ability to encode and retrieve past experiences in rich spatial contextual detail (episodic memory) are apparent by midlife – a time when most females experience spontaneous menopause. Yet, little is known about how menopause status affects episodic memory-related brain activity at encoding and retrieval in middle-aged pre- and post-menopausal females, and whether any observed group differences in brain activity and memory performance correlate with chronological age within group. We conducted an event-related task fMRI study of episodic memory for spatial context to address this knowledge gap. Multivariate behavioral partial least squares (PLS) was used to investigate how chronological age and retrieval accuracy correlated with brain activity in 31 premenopausal (age range: 39.55 – 53.30 yrs, *M_age_* = 44.28 yrs, *SD_age_* = 3.12 yrs) and 41 postmenopausal females (age range: 46.70 to 65.14 yrs, *M_age_* = 57.56 yrs, *SD_age_* = 3.93 yrs). We found that postmenopausal status, and advanced age within post-menopause, was associated with lower spatial context memory. The fMRI analysis showed that only in postmenopausal females, advanced age was correlated with decreased activity in occipitotemporal, parahippocampal, and inferior parietal during encoding and retrieval, and poorer spatial context memory performance. In contrast, only premenopausal females exhibited an overlap in encoding and retrieval activity in angular gyrus, midline cortical regions, and prefrontal cortex, which correlated with better spatial context retrieval accuracy. These results highlight how menopause status and chronological age, nested within menopause group, affect episodic memory and its neural correlates at midlife.

**Significance Statement:** This is the first fMRI study to examine how pre- and post-menopause status affects the neural correlates of episodic memory encoding and retrieval, and how chronological age contributes to any observed group similarities and differences. We found that both menopause status (endocrine age) and chronological age affect spatial context memory and its neural correlates. Menopause status directly affected the direction of age- and performance-related correlations with brain activity in inferior parietal, parahippocampal and occipitotemporal cortices across encoding and retrieval. Moreover, we found that only premenopausal females exhibited cortical reinstatement of encoding-related activity in midline cortical, prefrontal, and angular gyrus, at retrieval. This suggests that spatial context memory abilities may rely on distinct brain systems at pre-compared to post-menopause.

## Introduction

Aging is associated with declines in the ability to encode and consciously recollect past events in rich contextual detail (episodic memory; Tulving, 1972) and related brain activation differences in the medial temporal lobe (MTL), occipitotemporal cortex, prefrontal cortex (PFC), inferior parietal cortex, and midline cortical regions (Maillet & Rajah, 2013, 2014a; Persson et al., 2011; Rajah, 2015; Spaniol & Grady, 2010). Importantly, there is growing evidence that age-related episodic memory decline, as measured by object-location spatial context memory tasks, arises at midlife (Cansino et al., 2012) and is associated with altered PFC and occipitotemporal activity (Kwon et al., 2016). Midlife is also the time that most females with ovaries experience natural/spontaneous menopause and transition from being in pre-menopause to post-menopause (Harlow et al., 2012). Some postmenopausal females report cognitive issues, including brain fog and forgetfulness (Greendale et al., 2020). This raises the possibility that some of the prior reports of chronological age effects on memory and brain function at midlife may have been driven in part by postmenopausal females. Therefore, there is a need to bridge results from the cognitive neuroscience of aging and memory field with those from the menopause and endocrine aging field to advance our understanding how menopause status affects episodic memory and related brain function at midlife, and if any observed effects correlate with the chronological age of females at post-, compared to pre-, menopause (Taylor et al., 2019).

However, to our knowledge only one event-related task fMRI study has examined the effect of menopause status on brain activity during the performance of a verbal episodic encoding task (Jacobs et al., 2016). They found that postmenopausal status was associated with less activity in the hippocampus compared to premenopausal status. However, there was no behavioral effect of menopause status on verbal episodic retrieval, suggesting that the observed effects may not reflect the neural basis of memory disturbances experienced by some females at menopause (Greendale et al, 2020). Also, in this study the chronological age of pre- and post-menopausal females was matched to control for age effects. Thus, it remains unclear whether the observed menopause-related differences in brain activity differ with chronological age within pre- and/or post-menopause groups. Given that past work has suggested that brain and cognitive dysfunction at menopause is transient (Greendale et al., 2009), it is important to clarify whether and how chronological age affects memory and brain differences at pre, compared to post-menopause. Such information can help determine: i) what type of intervention may benefit females experiencing memory decline – ones that target endocrine function at menopause (e.g. hormone replacement therapies), if effects are stable; or ones that target both endocrine function and senescence, if effects vary with age at post-menopause (Doty et al., 2015); and ii) the timing and duration of these interventions. Finally, no study to our knowledge has examined how menopause status affects episodic retrieval-related brain activity, which highlights the paucity of knowledge about midlife brain health and episodic memory function in cognitively unimpaired females.

In the current task fMRI study, we investigate whether and how menopause status affects episodic memory and related brain function during encoding *and* retrieval in a broad sample of middle-aged premenopausal and postmenopausal females. We employ a face-location spatial context task fMRI paradigm that can detect episodic memory decline at midlife (Kwon et al., 2016). Performance on such tasks is correlated with the structural integrity and function of hippocampus (Beer et al., 2018; Snytte et al., 2020). Thus, we hypothesized that this paradigm would be sensitive for detecting post-menopause declines in episodic memory and associated differences in brain function. We also examine how chronological age, nested within menopause status, correlates with memory-related brain activity to determine if chronological aging contributes to any observed group similarities and/or differences. We hypothesize that there will be differences in chronological age affects on hippocampus, PFC and occipitotemporal cortex activity at encoding and retrieval based on menopause status.

## Materials and Methods

### Participants

Ninety-six middle-aged (39.5 – 65 years of age) females with ovaries participated in this study. Participants were categorized as being premenopausal (Pre-Meno) or postmenopausal (Post-Meno) based on the Stages of Reproductive Aging Workshop + 10 (STRAW +10) criteria (Harlow et al., 2012). Based on these criteria there were 33 Pre-Meno participants (Age range: 39.55 – 53.30 yrs, *M_age_* = 44.41, *SD_age_* = 3.12, *M_EDU_* = 16.44, Ethnicity: 81.8% White, 9.1% Latin American, 3.0% Black, 3.0% Chinese, 3.0% South Asian) and 63 Post-Meno participants (Age range: 46.70 to 65.14 yrs, *M_age_* = 57.05, *SD_age_* = 3.93, *M_EDU_* = 15.39, Ethnicity: 87.3% White, 3.2% Black, 3.2% Indigenous, 1.6% Chinese, 1.6% South Asian, 3.2% Middle Eastern and North African). This study was approved by the research ethics board of the Montreal West Island Integrated University Health and Social Services Centre – subcommittee for mental health and neuroscience, and all participants signed informed consent forms.

### Behavioral Methods

#### Enrollment

Participants were recruited via advertisements posted on websites and around the Montreal community. Participants first signed an online consent form and then proceeded to fill out an online questionnaire that collected demographics, medical history, reproductive history, and education information. This information was used to screen participants for inclusion/exclusion in the study. Individuals who met the inclusion/exclusion criteria listed below were invited to Behavioral Session 1 at the Cerebral Imaging Center (CIC) at the Douglas Mental Health University Institute (DMHUI).

Inclusion criteria: a minimum of high school education, willingness to provide a blood sample, and in general good health. Participants were excluded if they reported any of the following in their medical history: current use of hormone replacement therapy, bilateral oophorectomy, untreated cataract/glaucoma/age-related maculopathy, uncontrolled hypertension, untreated high cholesterol, diabetes, history of estrogen-related cancers, chemotherapy, neurological diseases or history of serious head injury, history of major psychiatric disorders, claustrophobia, history of a substance use disorder, or currently smoking >40 cigarettes per day. Participants were also excluded if they did not meet the requirements for magnetic resonance imaging (MRI) safety (e.g., implanted pacemaker). Females who were pregnant, perimenopausal or whose menopausal status was indeterminate, based on self-report and hormones, were excluded from the study.

#### Behavioral Session

After obtaining consent, participants filled out the MRI safety questionnaire. Next, they completed a battery of standardized psychiatric and neuropsychological assessments which included: the Mini-International Neuropsychiatric Interview (M.I.N.I; Sheehan et al., 1998), exclusion criteria was indications of undiagnosed psychiatric illness; the Beck Depression Inventory II (BDI-II; Beck et al., 1997), inclusion cut-off ≤ 19; the Mini Mental State Examination (MMSE; Folstein et al., 1975), inclusion cut-off ≥ 27. Participants then donated blood samples for hormonal assays to confirm self-reported menopausal status and performed a practice version of the spatial context memory task in a mock MRI scanner to familiarize them with the task and ensure they felt comfortable enough to partake in the real MRI scan. Participants who met the neuropsychological inclusion criteria stated above, were Pre-Meno or Post-Meno based on the STRAW+10 criteria and hormones (Harlow et al., 2012), and were able to perform the practice fMRI task, were invited to a second visit to undergo MRI scans at the CIC, DMHUI.

#### Endocrine Assessments

Estradiol-17β (E2), Follicle stimulating hormone (FSH), Luteinizing Hormone (LH) and Progesterone (P) levels were assessed to corroborate menopausal staging based on menstrual cycling and STRAW+10. Endocrine measurements were assessed on plasma, derived from heparin blood collection tubes. Blood was drawn on non-fasting individuals by a certified research nurse at the DMHUI during session 2 (MRI visit). Specimens were analyzed at the McGill University Health Centre (Glen site) Clinical Laboratories in Montreal. Endocrine Chemiluminescent immunoassays were performed on an Access Immunoassay System (Beckman Coulter), using the company’s reagents. All participants were categorized as Pre-Meno or Post-Meno based on the self-report and measured hormone levels for FSH as outline by STRAW+10 criteria. Females who were perimenopausal or whose menopausal status was indeterminate were excluded from the study.

#### Task fMRI Session

Upon arrival at the CIC, participants first took a pregnancy test. Participants who were not pregnant were then scanned while performing the spatial context memory task. Participants were asked to lie in a supine position in a 3T Siemens Prisma-Fit scanner, wearing a standard 32-channel head coil. High-resolution T1-weighted anatomical images were collected using a 3D Magnetization-Prepared Rapid Gradient-Echo (MP-RAGE) sequence (repetition time [TR] = 2300 ms, echo time [TE] = 2.36 ms, flip angle = 9°, 192 1 mm thick transverse slices, 1 mm × 1 mm × 1 mm voxels, field-of-view [FOV] = 256 mm^2^, acquisition time = 5:03 min) to use in the registration and normalization of the fMRI data. While performing the spatial context memory task, functional BOLD images were acquired using single shot T2*-weighted gradient echo-planar imaging (EPI) pulse sequence (TR = 2000 ms, TE = 30 ms, FOV = 256 mm^2^, matrix size = 64 × 64, in-plane resolution = 4 mm × 4 mm).

A mixed rapid event-related design was used for the task fMRI portion of the experiment (Dale, 1999). Participants were scanned during encoding and retrieval phases of easy and hard spatial context memory tasks. Task difficulty was manipulated by increasing encoding load (details below). A participant could perform up to four Easy and four Hard task runs (maximum eight runs total). Easy runs included 2 easy spatial context memory tasks (total run duration: 9:42 min), whereas hard runs only consisted of 1 experimental block (total run duration: 7:14 min). Run order was counterbalanced across all participants. Total length of the task fMRI session was approximately 1 hr 7 min and 44 sec.

The tasks were presented through E-Prime software (Psychology Software Tools, PA), back projected into the scanner bore, and made visible to participants via a mirror mounted within the head coil. Participants were provided with two 4-button inline fibre optic response boxes to respond throughout the task. During each run, response options and the corresponding buttons for each task were displayed at the bottom of the screen for clarity (see Figure 1 for a visual summary of the task fMRI protocol). Response accuracy and reaction time (RT, msec) was collected for all responses by E-Prime software.

**Figure 1:**
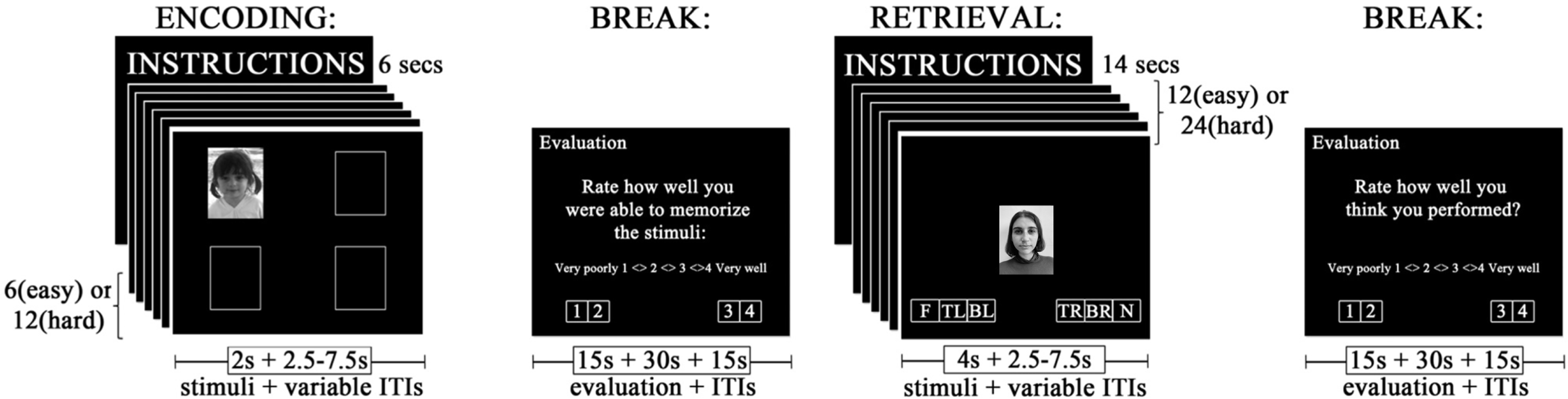
Task fMRI Paradigm Design. Research participants were asked to encode face-location associations and make a pleasant/neutral decision for face stimuli presented in one of four quadrants. There was a 1 minute interval between encoding and retrieval. During retrieval participants were presented with an equal number of previously encoded (“old”) and novel face stimuli in the centre of the screen and asked to make a 6-alterantive forced choice decision. Details about the paradigm are presented in the Materials & Methods section.

##### Encoding Phase

Participants were presented with instructions for the first six seconds of each encoding phase. Participants were informed that they would see a series of black and white photographs of faces that varied in age (from toddlers – older adults). They were instructed to memorize each face and its’ spatial location and to rate each face as either pleasant or neutral by pressing the corresponding button on the provided response box. The pleasantness rating was added to ensure participants deeply encoded the stimuli (Bernstein et al., 2002). The stimulus set has been described in detail in prior publications (Rajah et al., 2008). There were 50% girl/women, and 50% boy/men faces presented and the race and ethnicity of the faces presented were 83% White, 8.5% Black, 8.5% Other.

Following the instructions, participants were presented sequentially with either six or twelve face stimuli. Each face stimulus was presented one at a time (two sec/stimulus) in one of four quadrants on the computer screen (see Figure 1). The inter-trial interval (ITI) was varied (2.5 – 7.5 seconds; mean ITI = 4.17 seconds) to add jitter to fMRI data collection (Dale & Buckner, 1997). Conditions in which six faces were presented at encoding were considered the ‘Easy’, and those in which 12 faces were presented at encoding were considered ‘Hard’. Thus, the experiment included a task difficulty manipulation, which was based on the encoding load. This allowed us to assess within group and between group differences in performance as a function of increased task demands. Participants saw up to 48 faces during the easy encoding condition, and up to 48 faces presented during the hard encoding condition at the end of 4 easy runs. However, due to participants withdrawing participation and software errors, not all participants experienced all runs during the task fMRI session. The minimum number of easy runs experienced by any participant was two. Each easy run had two tasks, so the minimum number of easy tasks experienced was to four. Thus, the minimum number of faces encoded under ‘Easy’ condition was 4 tasks X 6 encoding stimuli = 24. The minimum number of hard runs experienced by any participant was three. Each Hard run had 1 Hard task. Thus, the minimum number of Hard encoding events experienced by a participant was 36. At the end of the encoding phase, participants rated their confidence on how well they performed the task on a four-point scale from very poorly (1) to very well (4) during a 60 second break.

##### Retrieval Phase

Participants were presented with instructions for fourteen seconds. They were informed that they would be shown a series of previously encoded (old) and novel (new) photographs of faces, presented to them one at a time. They were instructed to respond to each face by pressing one of six buttons corresponding to the following retrieval responses: (1) N – The face is NEW; (2) F – The face is FAMILIAR but I don’t remember its location; (3) TL – I remember the face and it was previously on the TOP LEFT; (4) BL – I remember the face and it was previously on the BOTTOM LEFT; (5) TR – I remember the face and it was previously on the TOP RIGHT; (6) BR – I remember the face and it was previously on the BOTTOM RIGHT. Participants were instructed not to guess and only respond (3) to (6) if they clearly recollected the face and its location.

Following the instructions participants saw a series of either 12 (Easy condition) or 24 (Hard condition) faces, presented one at a time, per task. Each face was presented for four seconds in the center of the screen. There was a variable ITI (2.5 – 7.5 s, mean ITI = 4.17 s). Participant saw up to a total of 96 faces (48 old, 48 new) at retrieval following 4 Easy and 4 Hard runs/conditions, respectively. The minimum number of Easy runs experienced by any participant was two. Each Easy run had two tasks, so the minimum number of Easy tasks experienced was to four. Thus, the minimum number of retrieval events under ‘Easy’ condition was 4 tasks X 12 encoding stimuli = 48 (24 old, 24 new). The minimum number of Hard runs experienced by any participant was three. Each Hard run had 1 hard task. Thus, the minimum number of Hard encoding events experienced by a participant was: 3 X 24 = 72 (36 old, 36 new). At the end of the retrieval phase participants were given 60 seconds to rate how well they thought they performed on a four-point scale from very poorly (1) to very well (4).

### Data analysis

#### Behavioral Analyses

Mean accuracy (% correct) and reaction time (msec) were calculated for each participant for the following retrieval response types within easy and hard tasks:

1. Correct Spatial Context Retrieval (CS) easy and hard: total number of *correct associative retrieval trials* (i.e., 3, 4, 5, 6 button presses made for previously seen faces, see Figure 1) divided by the total number of previously seen faces presented at retrieval.
2. Correct Recognition (Recog) easy and hard: total number of *correct retrieval* trials for recognition without memory for location (i.e. 2 responses for previously seen faces) divided by the total number of previously seen faces presented at retrieval.

### PLS Regression for Accuracy and RT

To examine the contributions of Menopause Group and Age (years) to variability in Recog and CS on Easy and Hard versions of tasks, as well as RT values for these judgments, we conducted PLS regression analyses. Specifically, we ran multi-response PLS regressions in *R* (version 4.1.3) using the *plsreg2* function from the *plsdepot* package (version 0.2.0; Sanchez, 2023). We selected this analytical method as it is capable of handling multicollinearity in predictor variables (as is the case for Menopause Group and Age) by creating a set of uncorrelated components to use in the model, and it can facilitate the inclusion of multiple response variables in one model (Abdi, 2010; Wold et al., 2001). The aim of PLS regression is to predict Y (i.e., the outcome variables) from X (i.e., the predictor variables) and describe their common structure. To achieve this, X and Y are first decomposed to form a set of latent variables (LVs) that maximally explain the shared variance (i.e., the covariance) between the original variables. The ability of a given LV to predict Y is then typically assessed via cross-validation, a process which produces Q^2^ values. LVs were considered significant and retained if Q^2^ ≥ 0.0975 (Abdi, 2010). For all such LVs, the variance explained (*R2*) in both X and Y is reported.

Although PLS regression is capable of handling cases involving highly collinear predictor variables, this type of analysis does not provide insight as to whether age is differentially related to spatial context retrieval accuracy at specific stages of menopause. That is, this method does not allow us to examine within group age effects. To address this, we conducted linear mixed-effects models for Pre-Meno and Post-Meno females in R using the lme4 package (version 1.1-29; Bates et al., 2015). Task Difficulty (Easy, Hard) and Age (Years) were entered as fixed effects, as was their interaction. Task Difficulty was coded using deviation coding and age was standardized. Both models included a random by-participant intercept. Statistical significance of fixed effects was determined via Satterthwaite approximations, implemented by the lmerTest package (version 3.1-3; Kuznetsova et al., 2017).

To assess whether effects of age provided a better fitting model, nested hierarchical linear mixed-effect models were fitted (with and without age as a predictor) and were then compared on values assessing goodness-of-fit (χ2) and model parsimony (AIC, Akaike Information Criterion; with ΔAIC ≥ 2 as the threshold for model selection).

### Task fMRI Analyses

#### Image Preprocessing

The DICOM files were first converted to NIFTI format and organized using the Brain Imaging Data Structure (BIDS; Gorgolewski et al., 2016). Volumes collected during the first 10 seconds of scanning were removed to ensure that magnetization had stabilized. Preprocessing was then conducted using fMRIprep version 20.2 (Esteban et al., 2019). FMRIprep is a robust preprocessing workflow in Python version 3.0 that implements tools from various software packages including Advanced Normalization Tools version 2.3.3 (ANTs), FMRIB Software Library version 5.0.9 (FSL), and FreeSurfer 6.0.1. <u>The following preprocessing description was generated by fMRIprep</u> (Esteban et al., 2019):

For each of the 8 BOLD runs found per participant (across all tasks and sessions), the following preprocessing was performed. First, a reference volume and its skull-stripped version were generated using a custom methodology of fMRIPrep. A B0-nonuniformity map (or fieldmap) was estimated based on a phase-difference map calculated with a dual-echo GRE (gradient-recalled echo) sequence, processed with a custom workflow of SDCFlows inspired by the epidewarp.fsl script and further improvements in HCP Pipelines (Glasser et al., 2013). The fieldmap was then co-registered to the target EPI reference run and converted to a displacements field map (amenable to registration tools such as ANTs) with FSL’s fugue and other SDCflows tools. Based on the estimated susceptibility distortion, a corrected EPI reference was calculated for a more accurate co-registration with the anatomical reference. The BOLD reference was then co-registered to the T1w reference using bbregister (FreeSurfer) which implements boundary-based registration (Greve & Fischl, 2009). Co-registration was configured with six degrees of freedom. Head-motion parameters with respect to the BOLD reference (transformation matrices, and six corresponding rotation and translation parameters) are estimated before any spatiotemporal filtering using mcflirt (FSL 5.0.9;Jenkinson et al., 2002). The BOLD time-series were resampled onto their original, native space by applying a single, composite transform to correct for head-motion and susceptibility distortions. These resampled BOLD time-series will be referred to as preprocessed BOLD in original space, or just preprocessed BOLD. The BOLD time-series were resampled into standard space, generating a preprocessed BOLD run in MNI152NLin2009cAsym space. First, a reference volume and its skull-stripped version were generated using a custom methodology of fMRIPrep. Several confounding time-series were calculated based on the preprocessed BOLD: framewise displacement (FD), DVARS and three region-wise global signals. FD was computed using two formulations following Power (absolute sum of relative motions, (Power et al., 2014)) and Jenkinson (relative root mean square displacement between affines; Jenkinson et al., 2002). FD and DVARS are calculated for each functional run, both using their implementations in Nipype (following the definitions by Power et al. 2014). Three global signals are extracted within the CSF, the WM, and the whole-brain masks. Additionally, a set of physiological regressors were extracted to allow for component-based noise correction (CompCor; Behzadi et al., 2007).

Principal components are estimated after high-pass filtering the preprocessed BOLD time-series (using a discrete cosine filter with 128s cut-off) for the two CompCor variants: temporal (tCompCor) and anatomical (aCompCor). tCompCor components are then calculated from the top 2% variable voxels within the brain mask. For aCompCor, three probabilistic masks (CSF, WM and combined CSF+WM) are generated in anatomical space. The implementation differs from that of (Behzadi et al., 2007) in that instead of eroding the masks by 2 pixels on BOLD space, the aCompCor masks are subtracted from a mask of pixels that likely contain a volume fraction of GM. This mask is obtained by dilating a GM mask extracted from the FreeSurfer’s aseg segmentation, and it ensures components are not extracted from voxels containing a minimal fraction of GM. Finally, these masks are resampled into BOLD space and binarized by thresholding at 0.99 (as in the original implementation). Components are also calculated separately within the WM and CSF masks. For each CompCor decomposition, the k components with the largest singular values are retained, such that the retained components’ time series are sufficient to explain 50 percent of variance across the nuisance mask (CSF, WM, combined, or temporal). The remaining components are dropped from consideration. The head-motion estimates calculated in the correction step were also placed within the corresponding confounds file. The confound time series derived from head motion estimates and global signals were expanded with the inclusion of temporal derivatives and quadratic terms for each (Satterthwaite et al., 2013). Frames that exceeded a threshold of 0.5 mm FD or 1.5 standardised DVARS were annotated as motion outliers. All resamplings can be performed with a single interpolation step by composing all the pertinent transformations (i.e., head-motion transform matrices, susceptibility distortion correction when available, and co-registrations to anatomical and output spaces). Gridded (volumetric) resamplings were performed using antsApplyTransforms (ANTs), configured with Lanczos interpolation to minimize the smoothing effects of other kernels. Non-gridded (surface) resamplings were performed using mri_vol2surf (FreeSurfer).

In addition to fMriprep, further preprocessing steps were carried out using custom code in Python 3.0 and Nilearn libraries (Abraham et al., 2014). If a participant had more than 15% of all their scans showing motion greater than 1 mm FD, they were removed from the analysis. For participants with motion <15% across all scans, the normalized scans from fMRIprep were scrubbed for motion artefacts as follows: a)when less than 15% of scans across all runs were above the 1mm cut-off, and only one or two consecutive volumes exceeded this cut-off, the volume was replaced by the average of the previous and the subsequent volume in time (e.g., a scan T_i that had an FD > 1mm was replaced by the mean/avg of scans T_(i-1) and T_(i+1)), b) when less than 15% of scans across all runs exceeded 1mm FD and there were 3 or more consecutive volumes that exceeded 1mm FD, then onsets including those volumes were excluded from the PLS analysis. The scans were then smoothed using a Gaussian filter (FWHM=6mm). Finally, confounds such as white matter, CSF and 6 motion parameters were regressed out from all the runs. The average percent (%) of total scans scrubbed post-processing was - 0.52% (1.04). The maximum % of scans scrubbed within a participant was 6.68%. The average number of onsets removed/censored - 0.35 (1.08). There were only 9 participants for whom onsets were removed. Maximum was 6 onsets excluded for a given participant.

#### Behavior PLS Analysis

A multivariate between-group behavior partial least squares (B-PLS; McIntosh & Lobaugh, 2004) analysis was conducted via open source PLSGUI software (https://www.rotman-baycrest.on.ca/index.php?section=345) in MATLAB (R2022a; Mathworks, Inc., Natick, MA). Partial least squares analysis was chosen because it is a powerful method that can be used to objectively assess spatiotemporal functional patterns in neuroimaging datasets. Moreover, between group B-PLS directly assesses the correlation between brain activity and different demographic and/or behavioral measures within group, i.e., age and memory performance (McIntosh et al., 2004). When conducting PLS the fMRI and behavioral data are stored in two separate matrices. The fMRI data matrix was organized with rows representing brain activity for each event-type of experimental interest nested within participants, and participants nested within Pre-Meno females and Post-Meno females groups. In the current analysis the rows included activity for the following four event types of interest, nested within participant, nested within group: all encoding events during easy conditions (EncEasy), all encoding events during hard conditions (EncHard), all retrieval events during easy conditions (RetEasy), all retrieval events during hard conditions (RetHard). The columns of the fMRI data matrix contain mean-centred BOLD activity of all brain voxels at event onset and the subsequent 7 scans/lags, wherein each lag is equivalent to the T2*-weighted gradient EPI sequence’s TR or 2000 msec; thus capturing 16 sec of whole-brain activity following every event onset. PLS averages the number of events for each event-type/condition by lag. As stated above, the minimum number of encoding events experienced by a participant during easy conditions was 24, and during hard condition it was 36. The minimum number of easy retrieval events experienced by a participant during easy conditions was 48 (24 old, 24 new), and during hard conditions it was 72 (36 old, 36 new).

The behavioral data were stored in a separate matrix. The rows of the behaviour matrix are stacked in the same order as the fMRI data matrix The columns of the behavioral matrix included vectors of participants’ behavioral data for: (1) standardized age (2) standardized mean spatial context accuracy and (3) standardized mean recognition accuracy. The behavior matrix was cross correlated with the fMRI data matrix and underwent singular value decomposition (SVD). SVD yields a series of orthogonal latent variables (LVs) that is equal to the number of groups-by-event-type/condition (2 groups × 4 event-type/conditions = 8 LVs). Each LV is composed of a singular value, a singular image, and a correlation profile. The singular value represents the proportion of covariance accounted for by the LV. The singular image includes positive and/or negative saliences which establish the weighted contributions of each voxel at a given time lag, producing a spatiotemporal pattern of whole-brain activity. Time lags are the number of TRs after the stimulus of interest is presented. The correlation profile portrays the association between the pattern of brain activity from the singular image with the behavioral measures. In interpreting the correlation profiles, it is important to note that PLS results are symmetrical. Such that, negative correlations between a behavioral vector (age, performance) and a negative salience brain regions is symmetrical with more activity in these negative salience regions being positively correlated with these same behavioral vectors. The opposite symmetry can be stated for positive correlations with positive salience regions.

Permutation testing was conducted on the LVs to establish significance (*p* < 0.05; McIntosh et al., 2004). The permutation test involved sampling without replacement to reassign links between subjects’ behavioural vector measures and event/condition within subject. For each permuted iteration a PLS was recalculated, and the probability that the permuted singular values exceeded the observed singular value for the original LV was used to assess significance at *p* < 0.05 (McIntosh et al., 2004). To identify stable voxels that consistently contributed to the correlation profile within each LV, the standard errors of the voxel saliences for each LV were estimated via 500 bootstraps, sampling subjects with replacement while maintaining the order of event types for all subjects. For each voxel, a value similar to a z-score known as the bootstrap ratio (BSR) was computed, reflecting the ratio of the original voxel salience to the estimated standard error for that voxel. Voxels with BSR values of ± 3.28 (equivalent to *p* < 0.001) and a minimum spatial extent = 10 contiguous voxels, were retained and highlighted in the singular image. BSR values reflect the stability of voxel saliences. A voxel salience whose value is dependent on the observations in the sample is less precise than one that remains stable regardless of the samples chosen (McIntosh & Lobaugh, 2004). The bootstrapping results were used to calculate 95% confidence intervals (CI) for the correlations between event-related brain activity and the three behavioral vectors of: chronological age, spatial context retrieval accuracy, and recognition accuracy during EncEasy, EncHard, RetEasy, RetHard conditions. If the CI crosses zero, the effect is not significant for that group and condition. In this B-PLS analysis a group difference in the correlation between chronological age and memory-related brain activity would manifest as either the correlations between age and memory-related brain activity being in opposite directions between Pre-Meno and Post-Meno; or, as one group showing a correlation between age and memory-related brain activity and not the other. Group similarities in age differences in memory-related brain activity would present as both groups exhibiting the same direction of correlation between age and brain activity.

Temporal brain scores were calculated to obtain time lags with the strongest correlation profile for significant LVs from each PLS analysis. Temporal brain scores reveal the strength of the pattern of brain activity for each participant at each time lag. We retained only peak coordinates from time lags 3 to 5, which exhibited maximal effects related to the PLS effect. Identified peaks were converted to Talairach space using the icbm2tal transform (Lancaster et al., 2007), implemented in GingerAle 2.3(Eickhoff et al., 2009). Relevant Brodmann areas were established using the Talairach and Tournoux atlas (Talairach & Tournoux, 1988) and confirmed in FSL. Peak coordinates from the cerebellum and brainstem were not included as the fMRI acquisition in these regions was incomplete.

#### Post-hoc analyses

Post-hoc between group repeated measure MANOVAs were conducted for each significant LV to determine if there were significant group differences in LV brain scores based on female’s current antidepressant use, HBC use and diagnosis of PCOS. Significance was assessed at *p* < 0.05 corrected.

### Data and code availability

Code (MATLAB, R) used for the present study is made publicly available on our Lab GitHub page (https://github.com/RajahLab/BHAMM_TaskfMRI_scripts). Readers seeking access to data should email the Principal Investigator of the Brain Health at Midlife and Menopause (BHAMM) Study, Professor Maria Natasha Rajah, for further information.

## Results

### Behavioral results

A total of 24 participants were excluded from behavioral and fMRI analyses: 11 Post-Meno were excluded for being on HRT (9) or HBC (2) for hormonal regulation, 1 participant due to incidental finding, 1 participant due to scanner technical issues, 2 participants due to excessive movement (> 15% of all scans with FD > 1mm) during MRI scanning, 8 participants for being behavioral outliers and/or task non-compliant, and 1 participant was identified as a PLS brain scores-behaviour correlation outlier. A behavioral outlier was defined as an individual that had a Cook’s D value which was 3 stand deviations from the group mean on 3 or more response types. A non-compliant responder was defined as someone who performed poorer than chance on the correct rejections and/or had zero spatial context retrieval responses.

Demographic and fMRI behavioral data separated by reproductive stage for the final sample (*N* = 72) are reported in Table 1. The age range for the 31 Pre-Meno females was 39.55 to 53.30 years, with an education range of 12 – 20 years. The age range for the 41 Post-Meno females was 52.30 to 65.14 years, with an education range of 11 to 20 years. Chi-square tests were conducted to determine if there were group differences in the proportion of antidepressant medication use, use of hormonal birth control (HBC, i.e., oral contraceptive, patch, intrauterine devices), or diagnosis of polycystic ovary syndrome (PCOS). There was a significant group difference in HBC use (*p* = 0.018), with Pre-Meno having 12.0% HBC users for (*n* = 4) and Post-Meno females having 0%. We did not exclude the 4 Pre-Meno HBC users as their reproductive health data indicated it was being used for contraceptive purposes, and they were not identified as outliers in the behavioral or fMRI analyses. One-way ANOVAs were conducted to compare Pre-Meno and Post-Meno females on demographic variables (age, education, handedness, body mass index [BMI], BDI-II scores), blood-based measures of sex hormone levels (E2, FSH, LH, P), and task fMRI behavioral data. There was a significant group difference in age, and hormone levels. Post-Meno females were significantly older than Pre-Meno females. Post-Meno females had significantly higher FSH, LH levels, and lower E2 and P, levels compared to Pre-Meno females, consistent with their menopause staging. Detailed analysis of the task fMRI analysis data is presented below.

**Table 1:** Demographic Information and fMRI Behavioural Measures by Reproductive Group. Handedness assessed using Edinburgh Handedness Inventory. Antidepressant use based on self-report. BMI = body mass index; BDI-II = Beck Depression Inventory-II; BAI = Beck Anxiety Inventory; CVLT = California Verbal Learning Test; LFR = Long-form Free Recall; LCR = Long-form Cued Recall; RG = Recognition; HBC = Hormone-Based Contraceptive; HRT = Hormone Replacement Therapy (% on each medication); PCOS (%) = Polycystic ovarian syndrome (% with PCOS diagnosis). Due to detection thresholds of the ELISA used, E2 was measurable in 12 out of 48 Post-Meno females who donated blood for hormonal measures, all other hormone measures in Post-Meno females are the average from 48. For Pre-Meno females all measures are based on 28 of 31 who donated blood for hormonal analysis. CS = Correct Spatial Context Retrieval; CR = Correct Rejection; ACC = Accuracy; RT = Reaction Time. Accuracy values are shown as proportion correct per task type with standard error (SE). Reaction time values are shown in milliseconds (msec) per task type. SPSS (Version 29, IBM Corporation) was used to conduct Chi-square tests for demographics, and one-way between-group ANOVAs for hormones, and fMRI Behavioral measures to test for group differences. For the fMRI Behavioral significance was assessed at p<0.05 corrected for 20 multiple comparisons (p<0.0025 uncorrected) for accuracy. For RT, measures are only possible for 7 of the response types and corrections were for 14 response types, p<0.0035 uncorrected or p<0.05 corrected.

| | Premenopausal | Postmenopausal | Effect size ($\eta^2$/Cramer's V) |
| --- | --- | --- | --- |
| Sample size (*n*) | 31 | 41 |  |
| Age (years)* | 44.28 (0.57) | 57.56 (0.56) | 0.793 |
| Education (years)<sup>§</sup> | 16.34 (0.40) | 15.21 (0.40) | 0.052 |
| BMI (kg/m<sup>2</sup>) | 25.14 (1.00) | 25.45 (0.63) | 0.001 |
| BDI-II | 4.10 (0.71) | 6.10 (0.87) | 0.040 |
| Handedness (% right) | 90.32 (2.56) | 87.07 (4.80) | 0.363 |
| Antidepressant (#) | 7 | 8 | 0.037 |
| HBC (#)* | 4 | 0 | 0.279 |
| PCOS (#) | 3 | 2 | 0.096 |
| E2 (pmol/L)* | 355.61 (42.44) | 35.23 (3.10) | 0.546 |
| FSH (IU/L)* | 9.19 (1.68) | 85.69 (4.45) | 0.759 |
| LH (IU/L)* | 6.60 (1.05) | 34.22 (1.95) | 0.665 |
| P (nmol/L)* | 17.52 (3.49) | 1.54 (0.15) | 0.308 |
| % CS easy Accuracy* | 63.91 (3.61) | 44.78 (2.86) | 0.202 |
| % Recog easy Accuracy* | 13.17 (1.87) | 22.68 (2.71) | 0.094 |
| % CS hard Accuracy* | 43.08 (3.19) | 28.42 (2.25) | 0.176 |
| % Recog hard Accuracy* | 25.34 (2.50) | 33.30 (2.90) | 0.054 |
| CS easy RT (msec)* | 2429.81 (82.32) | 2757.01 (84.94) | 0.094 |
| Recog easy RT (msec) | 3011.28 (113.74) | 3047.63 (125.06) | 0.001 |
| CS hard RT (msec)* | 2595.57 (76.04) | 2936.40 (86.12) | 0.105 |
| Recog hard RT (msec) | 2889.18 (105.49) | 2762.19 (99.14) | 0.011 |

Note: Sample size and Mean (SE) for demographic, reproductive health, and task fMRI behavioural data. Significant p-value ( $p < 0.05$ ) for between-group analysis using two-sided Chi-square or one-way ANOVA marked with '\*'. <sup>§</sup>The one-way ANOVA for education was nearly significant with $p = 0.055$ .

#### Behavioral Analysis of Task fMRI Accuracy and RT data

The PLS regression analysis of accuracy data identified one LV with a Q^2^ value above threshold (Q^2^ = 0.1572). This LV accounted for 94.47% of the variance in X (i.e., Menopause Group, Age) and 17.81% of the variance in Y (i.e., CS Easy/Hard, Recog Easy/Hard). Figure 2a highlights the correlations between the original predictor/outcome variables and the LV. For the predictor variables, both Menopause Group and Age were negatively associated with the LV. For the outcome variables, Recog Easy and Recog Hard were negatively correlated with the LV (albeit relatively weakly), whereas CS Easy and CS Hard were positively correlated with the LV. These results indicate that the LV captured a pattern in which Menopause Group and Age were positively correlated with Recog but negatively correlated with CS, for both Easy and Hard conditions. This indicates that both predictor variables were associated with changes in spatial context memory performance. By contrast, the PLS regression analysis of RT data failed to identify an LV with a Q2 value above 0.0975 (Abdi, 2010).

**Figure 2:**
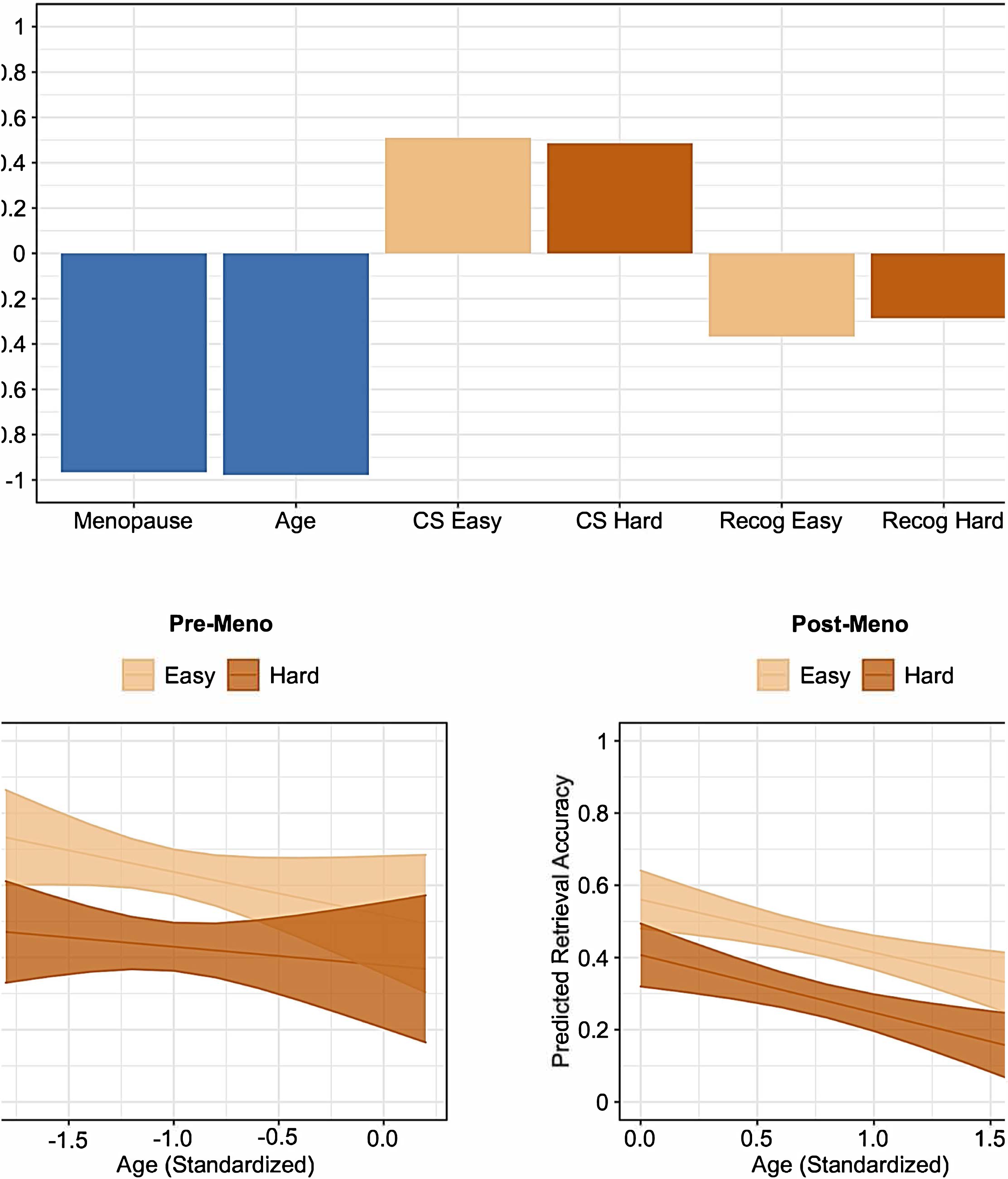
Behavioral Results. A) The correlation profile for the LV identified by PLS regression. The bars reflect the correlation between the original independent (blue)/dependent (orange) variables and the LV; B) Association between age (standardized) and spatial context retrieval accuracy as a function of task difficulty (easy, hard) in Pre-Meno females (left) and Post-Meno females (right). Lines of best fit and 95% confidence intervals are based on predicted values.

As a follow-up to this analysis, we conducted linear mixed-effects models to test the effect of Age and Task Difficulty (Easy, Hard) on CS within each group. For Pre-Meno females, we observed a statistically significant effect of Task Difficulty (F(1, 29) = 4.905, p = .035) but not Age (F(1, 29) = 1.275, p = .268). The effect of Task Difficulty was driven by lower CS on the Hard compared to the Easy task version. Moreover, there was no statistically significant interaction (F(1, 29) = 1.424, p = .242). For Post-Meno females, we observed statistically significant effects of Task Difficulty (F(1, 39) = 20.26, p < .001) and Age (F(1, 39) = 11.372, p = .002). These effects were driven by lower CS on the Hard compared to the Easy task version and with advanced age. Notably, no interaction effect was present (F(1, 39) = 0.125, p = .726). An overview is presented in Figure 2b. These results suggest that the effect of chronological age on spatial context retrieval accuracy is dependent on menopausal status, reflecting a more complex pattern than was evident in the PLS regression analysis.

Comparison of hierarchical linear mixed-effects models showed that, for Pre-Meno females, adding Age to a model with Task Difficulty did not provide a significantly better fitting or a more parsimonious model, χ2=2.8, p=.24, ΔAIC = -1. For Post-Meno females, adding Age to a model with Task Difficulty significantly improved model fit and parsimony, χ2=10.6, p=.005, ΔAIC = -7. These results suggest that adding chronological age as a predictor provides a better explanatory model of the variability in the dependent variables for Post-, but not Pre-Menopausal females.

#### Post-hoc Linear Regressions to Explore Education Effects

Given that there was a trend in group differences in education, with Pre-Meno females trending towards being more educated than Post-Meno females (*p* = 0.055), we conducted four linear regressions models to explore if Menopause Group (-1 Pre-Meno, +1 Post-Meno), standardized Education (zEDU), and the Menopause Group*zEDU interaction predicted CS easy, CS hard, Recog easy, and Recog hard. There was no significant Menopause Group*zEDU interaction observed in any of the models (range of p-values 0.30 – 0.99). There was a significant effect of zEDU for both Menopause Groups for CS hard accuracy (standardized β = 0.246, *t* = 2.17, *p* = 0.033).

### Task fMRI results

The B-PLS analysis was conducted to determine how menopause status affected correlations between memory performance and brain activity at encoding and retrieval; and how this varied with chronological age within group. Performance was assessed using both spatial context and recognition accuracy to explore if groups engaged similar or different brain regions to support these two different response-types. The B-PLS fMRI analysis identified two significant LVs. The post-hoc between menopause group repeated multivariate analysis of variance (MANOVA) to assess whether there was an effect of female’s history of antidepressant use, hormonal birth control (HBC) use, and diagnosis of Polycystic Ovary Syndrome (PCOS) on PLS brain scores identified no significant main effects or interactions for the two significant LVs (*p* > 0.05, LV1 and LV2).

#### LV 1 - Correlations between retrieval activity and memory performance in Pre-Meno females

LV 1 accounted for 14.57% of cross-block covariance and was significant at *p* < 0.001 based on permutation testing. Figure 3A presents the brain-behaviour correlation profile for LVI, with 95% confidence intervals (CI) for the correlation effects based on the bootstrapping, and summarizes how chronological age, spatial context retrieval accuracy, and recognition accuracy correlated with brain activity in regions identified in the singular image (Figure 3C) during EncEasy, EncHard, RetEasy, RetHard conditions. If the CI crosses zero, the effect is not significant for that group and condition. Figure 3B shows the scatterplots of brain scores by age and performance for Pre-Meno and Post-Meno females. Table 2 presents the local maxima for LV 1. This LV primarily identified negative salience brain regions which included: dorsolateral prefrontal cortex (DLPFC), ventrolateral PFC, angular gyrus, and midline cortical regions, i.e., medial PFC (mPFC), anterior cingulate, posterior cingulate and precuneus.

**Figure 3:**
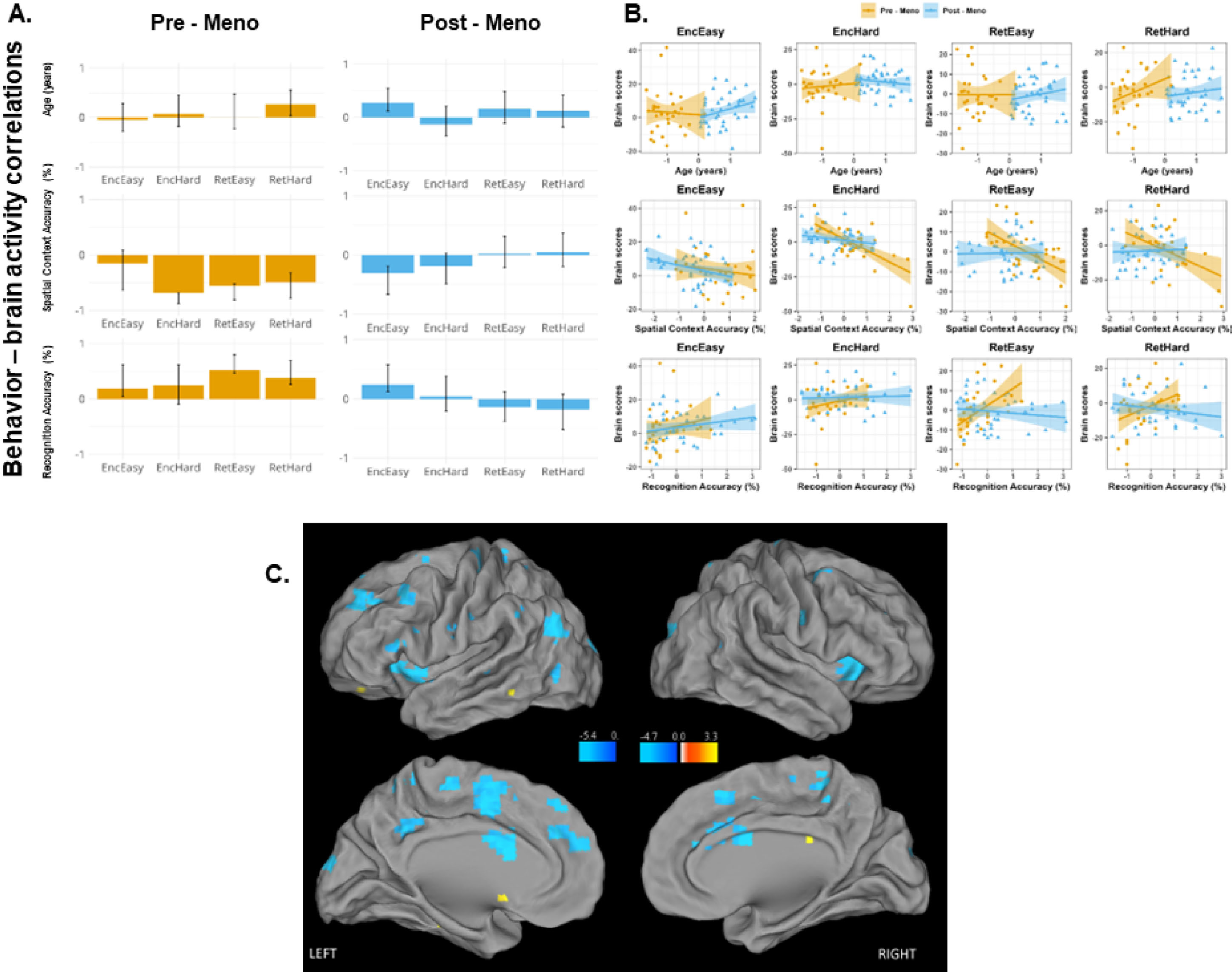
Brain-behavior PLS Results – LV 1. A) The brain-behaviour correlation profiles for LV1 depicting how chronological age, spatial context retrieval accuracy, and at item retrieval accuracy correlated with activity in regions identified in the singular image. The bars represent the correlation values between age, spatial context retrieval accuracy, and recognition accuracy and the brain activity pattern identified by the symmetrically paired singular image. The whiskers on the bars reflect the 95% confidence intervals (CI) based on the bootstrap results. Bars in which the whiskers cross zero indicate that correlation effect did not significantly contribute to the results; B) Presents scatterplots which depict the correlations between subjects’ brain scores and age, spatial context retrieval accuracy, and recognition accuracy for Pre-Meno (in yellow) and Post-Meno (in blue) females. The slope of these correlations, with 95% CI, represent the bar values and whiskers in (A), respectively; C) The singular image for LV1, threshold bootstrap ratio of ± 3.28, *p* < 0.001. Only negative salience brain regions were identified and are coloured in blue. The figure highlights regions identified between TR/Lags 3 – 5 based on the temporal brain scores. The scale colour codes effects according to the strength of BSR. Interpretation of this LV is presented in the Results section. The singular image is presented on template images of the lateral and medial surfaces of the left and right hemispheres of the brain using Caret software v5.65 (http://brainvis.wustl.edu/john/caret5).

**Table 2:**
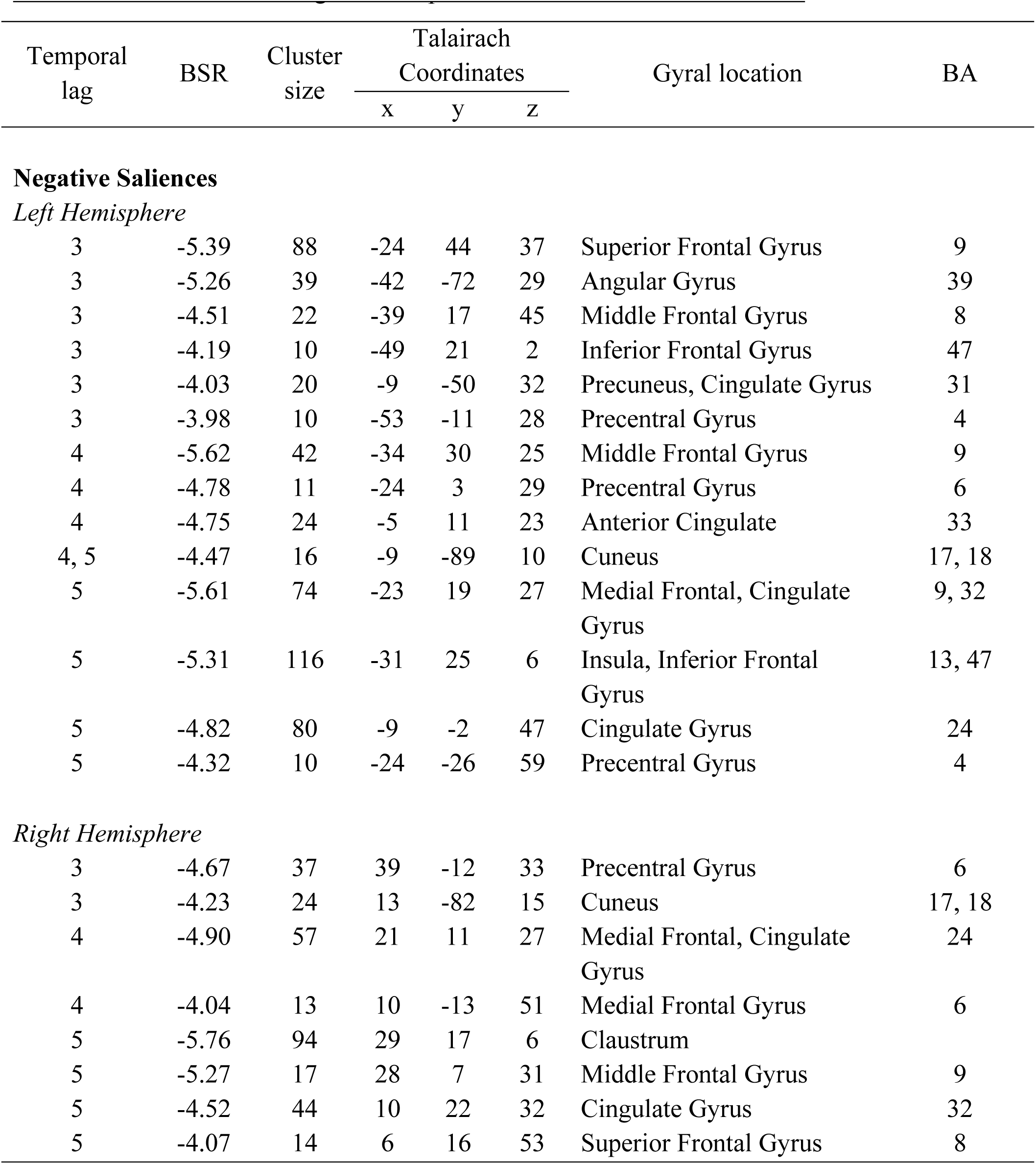
Local maxima of negative and positive saliences identified in LV 1. Temporal lag represents the time after event onset when a cluster of voxels showed an effect of interest. The temporal brain score identified lags 3 – 5 as maximally representing the LV effects identified by the PLS. The bootstrap ratio (BSR) threshold was set to ± > 3.28 (*p* < 0.001) and identified dominant and stable activation clusters. The spatial extent refers to the total number of voxels included in the voxel cluster (threshold = 10). The stereotaxic coordinates (measured in millimeters), gyral location and Brodmann areas (BAs) were determined by referring to Talaraich and Tournoux (1998). HEM = Cerebral hemisphere in which the activation of interest occurred.

Overall, Pre-Meno females accounted for LV 1 effects. In Pre-Meno females, increased activity in negative salience regions during EncHard, RetEasy, and RetHard conditions was correlated with better spatial context memory. Importantly, retrieval effects were only observed in Pre-Meno females. Pre-Meno females who exhibited more retrieval activity in negative salience regions also had higher spatial context memory accuracy and lower recognition accuracy across Easy and Hard conditions. In addition, in Pre- Meno females age was correlated with less retrieval activity in these brain regions during the RetHard conditions.

Although LV1 primarily reflected correlations between retrieval activity and memory performance in Pre-Meno females, Figure 3A and 3B indicate that both Pre-Meno and Post-Meno females who exhibited more activity in negative salience regions during encoding had greater subsequent spatial context memory accuracy, and lower subsequent recognition accuracy. As noted above, in Pre-Meno females this effect was only observed for the EncHard conditions, and in Post-Meno females this effect was only in the EncEasy conditions. Also, only in Post-Meno females this effect was correlated with age. Older Post-Meno females exhibited less activity in these regions during EncEasy and this was linked to lower subsequent spatial context accuracy and higher subsequent recognition for the Easy condition.

#### LV 2 - Menopause Group differences in age and performance related patterns of brain activity

LV2 (*p* = 0.013) accounted for 10.34% of cross-block covariance (see Figure 4, Table 3). This LV primarily identified negative salience brain regions that included: ventral occipitotemporal, bilateral parahippocampal gyrus and bilateral parietal cortices. The correlation profile and brain – behavior scatterplots (Figure 4A, 4B) indicate that LV 2 identified group differences in age-, and performance-related brain activity. Post-Meno females contributed more strongly to this LV. Specifically, Post-Meno females who exhibited more activity in negative salience brain regions during encoding and retrieval also had higher spatial context retrieval accuracy and lower recognition accuracy. However, with advanced age within Post-Meno females, there was less activity in negative salience regions, and thus lower spatial context accuracy and higher recognition accuracy.

**Figure 4:**
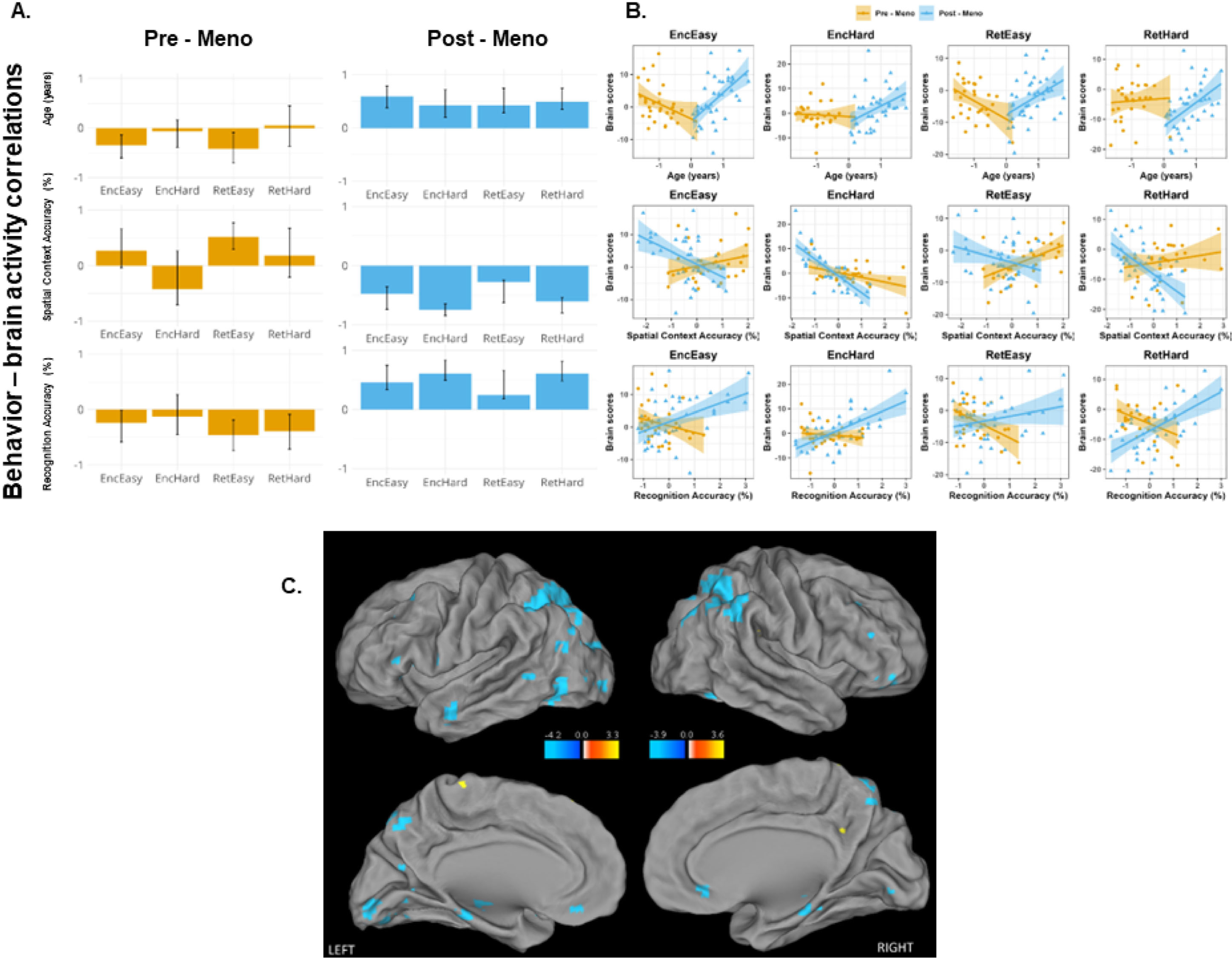
Brain-behavior PLS Results – LV 2. A) The brain-behaviour correlation profiles for LV 2 depicting how chronological age, spatial context retrieval accuracy, and at item retrieval accuracy correlated with activity in regions identified inf the singular image; B) Presents scatterplots which depict the correlations between subjects’ brain scores and age, spatial context retrieval accuracy, and recognition accuracy for Pre-Meno (in yellow) and Post-Meno (in blue) females. The slope of these correlations, with 95% CI, represent the bar values and whiskers in (A), respectively; C) The singular image for LV2, threshold bootstrap ratio of ± 3.28, *p* < 0.001. Only negative salience brain regions were identified and are coloured in blue. The figure highlights regions identified between TR/Lags 3 – 5 based on the temporal brain scores. The scale colour codes effects according to the strength of BSR. The singular image is presented on template images of the lateral and medial surfaces of the left and right hemispheres of the brain using Caret software v5.65 (http://brainvis.wustl.edu/john/caret5). Interpretation of this LV is presented in the Results section.

**Table 3:**
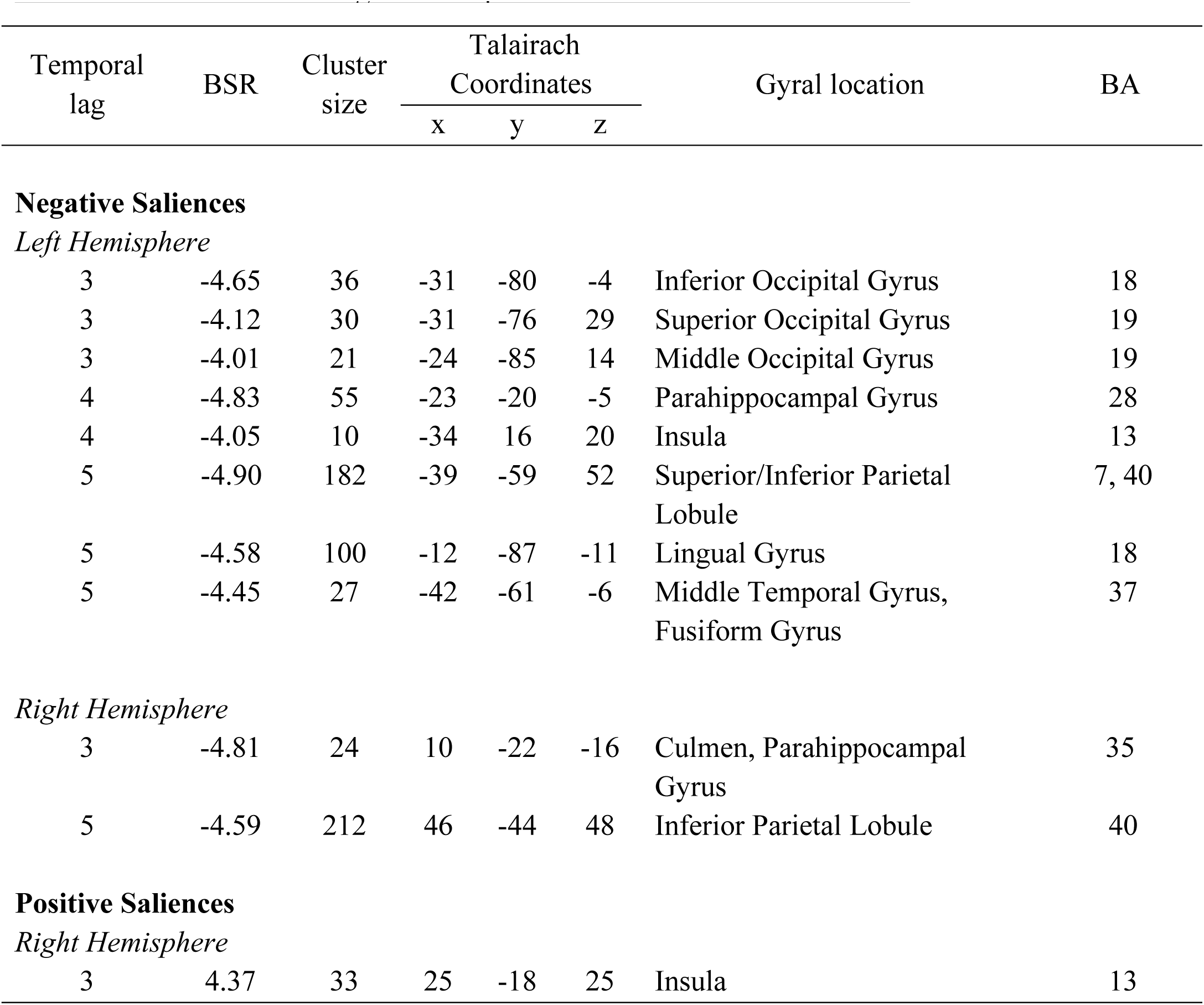
Local maxima of negative and positive saliences identified in LV 2. Temporal lag represents the time after event onset when a cluster of voxels showed an effect of interest. The temporal brain score identified lags 3 – 5 as maximally representing the LV effects identified by the PLS. The bootstrap ratio (BSR) threshold was set to ± > 3.28 (*p* < 0.001) and identified dominant and stable activation clusters. The spatial extent refers to the total number of voxels included in the voxel cluster (threshold = 10). The stereotaxic coordinates (measured in millimeters), gyral location and Brodmann areas (BAs) were determined by referring to Talaraich and Tournoux (1998). HEM = Cerebral hemisphere in which the activation of interest occurred.

The effects in Pre-Meno females were in the opposite direction to those observed in Post-Meno females and were more complex. Advanced age in Pre-Meno females was correlated with more activity in negative salience brain regions during Easy conditions only. Also, Pre-Meno females who exhibited less activity in these regions during RetEasy had lower spatial context retrieval accuracy and higher recognition accuracy. Similarly, Pre-Meno females who exhibited more activity in these regions during the RetHard conditions also had higher recognition accuracy during these conditions. Thus, LV2 identified Menopause Group differences in age and performance related patterns of brain activity.

## Discussion

In the current study we used multivariate behavioral and functional neuroimaging analysis methods to examine how the menopause status affected episodic memory for face-location association (spatial context memory) and its functional neural correlates, and how this correlated with age and memory performance within group. The behavioral PLS regression analysis indicated that both older age and Post-, compared to Pre-, menopause status was related to lower spatial context retrieval accuracy and higher recognition accuracy. This result is consistent with past work that has indicated that Post-Meno females perform poorer on associative memory tasks compared to age-matched Pre-Meno females (Rentz et al., 2017) and that context memory accuracy is lower at midlife compared to young adulthood (Kwon et al., 2016). When probed within groups, however, we found that advanced age was associated with poorer spatial context retrieval accuracy in Post-, but not Pre-, menopausal females. This suggests that the detrimental role of age on spatial context memory emerges in late-midlife, at a time when females with ovaries have already transitioned through menopause. Overall, our findings show that older Post-Meno females experience deficits in spatial contextual recollection, in the presence of intact familiarity-based item retrieval and/or item-specific recollection (Yonelinas et al., 1996).

The multivariate task fMRI analysis identified two significant LVs. The first LV (LV 1) identified both similarities and differences in the correlations between age, episodic retrieval accuracy, and brain activity during encoding and retrieval. LV 2 only identified group differences in the correlations between brain activity, age, and memory performance. Below we discuss our task fMRI findings in greater detail.

### Group similarities in subsequent memory effects

LV 1 identified significant correlations between encoding activity in lateral PFC, angular gyrus, and midline cortical regions, and subsequent memory performance in both Pre-Meno and Post-Meno females. Specifically, middle-aged females who exhibited more activity in these regions at encoding also exhibited better spatial context accuracy (and lower recognition accuracy). The midline cortical regions identified in this LV (i.e., medial PFC, cingulate and precuneus regions) are part of the default mode network (DMN (Biswal et al., 1997; Greicius et al., 2003; Power et al., 2010)). Prior episodic memory studies have found that greater activity in these regions is associated with successful episodic encoding and contextual recollection (de Chastelaine et al., 2016; Kim, 2010; Rugg & Vilberg, 2013). Similarly, increased activity in lateral PFC activity has been consistently observed during spatial context memory tasks at encoding and retrieval (Mitchell & Johnson, 2009; Rajah et al., 2008; Rajah et al., 2010), and increased left angular gyrus activity has been observed during associative object-context encoding and recollection (Bellana et al., 2023; Branzi et al., 2021; Maillet & Rajah, 2014b). In addition, Jacobs et al (2016) reported no significant group differences in lateral PFC activity during a verbal episodic encoding tasks between Pre-Meno and Post-Meno females, consistent with our current findings.

Interestingly, in the current study Post-Meno females exhibited subsequent memory effects in the aforementioned brain regions during the EncEasy condition, whereas Pre-Meno females exhibited these effects during the EncHard condition. Moreover, in Post-Meno females, the subsequent memory effects during EncEasy also correlated with chronological age. Older Post-Meno females exhibited less activity in the mentioned brain regions, and this was correlated with poorer subsequent spatial context memory. Given that the difficulty manipulation in the current study was one of encoding load, our results show that older Post-Meno females displayed a pattern of brain activity consistent with the prediction of the Compensation Related Utilization of Neural Circuits Hypothesis (CRUNCH): older Post-Meno females activated similar regions as younger Pre-Meno females, but at a lower levels of task demands, which may reflect compensation for neural inefficiencies (Cappell et al., 2010).

### Group differences in age and performance-related brain activity

LV1 and LV2 identified group differences in how chronological age affected memory-related brain activity and corroborates our hypothesis that at a neural level there are menopause-related differences in the effect of age on memory-related brain activity. For example, in LV 1, group differences were observed at retrieval. Pre-Meno females who exhibited more activity in lateral PFC, angular gyrus and midline cortical activity during easy and hard retrieval conditions also had higher spatial context memory accuracy and lower recognition accuracy. Therefore, only in Pre-Meno females was reinstatement of encoding-related activity in lateral PFC, angular gyrus and midline cortical regions at retrieval correlated with better spatial context memory. Moreover, age-related declines in this reinstatement during the more challenging RetHard conditions was detrimental to spatial context memory in only Pre-Meno females.

Past work has shown how reinstatement of encoding-related activity at retrieval is important for successfully remembering episodic memories (Gordon et al., 2013; Hill et al., 2021; Leiker & Johnson, 2015; Thakral et al., 2015). Different patterns of cortical reinstatement may support successful recollection and context retrieval, compared to successful recognition (Johnson et al., 2009). In a previous fMRI study Thakral et al (2015) reported that in young adults, cortical reinstatement of the same regions identified in LV1 (lateral PFC, angular gyrus, midline cortical regions) supported successful retrieval of context information. Johnson et al (2009) reported that reinstatement of posterior midline cortical activity was particularly relevant for successful recollection in a young adult sample. Therefore, our results indicate that middle-aged Pre-Meno females engage similar brain regions to young adults to support successful spatial context memory.

In contrast, middle-aged Post-Meno females did not show cortical reinstatement of encoding related activity in lateral PFC, angular gyrus and midline cortical at retrieval. Importantly, there was also no significant correlation between chronological age and LV 1 retrieval activity in Post-Meno females. However, LV2 results show that Post-Meno females can engage in cortical reinstatement; but the regions they reactivated at retrieval were distinct from those engaged by Pre-Meno females to support spatial context memory. Specifically, LV2 shows that Post-Meno females who exhibited greater activity in bilateral occipitotemporal, parahippocampal, and inferior parietal cortices at encoding *and* retrieval had higher spatial context memory accuracy and lower recognition memory accuracy. Interestingly, Johnson et al (2009) found that cortical reinstatement effects in occipitotemporal regions supported both detailed recollection and more general recognition memory. However, increased activity in parahippocampal and inferior parietal has explicitly been observed during spatial context memory tasks (Ankudowich et al., 2016; Diana et al., 2013; Hayes et al., 2007; Snytte et al., 2022). LV2 PLS results also indicated that older Post-Meno females exhibited lower levels of bilateral occipitotemporal, parahippocampal and inferior parietal cortex activity at encoding and retrieval, which was correlated with lower spatial context memory and higher recognition memory. Taken together, LV 1 and LV 2 results suggest that spatial context memory reductions in Post-Meno females compared to Pre-Meno females may in part be due to menopause/endocrine aging affecting the ability to reactivate lateral PFC, midline cortical and angular gyrus at retrieval (LV1) (Bellana et al., 2023; Morcom, 2014; Vilberg & Rugg, 2014; Wing et al., 2015); and, chronological age-related reductions in occipitotemporal, parahippocampal and inferior parietal activity at encoding and retrieval. These results highlight the combined influences of menopause and chronological aging on episodic memory and related brain function in middle-aged Post-Meno females.

LV 2 also showed that middle-aged Pre-Meno females exhibited correlations between age, performance, and brain activity in occipitotemporal, parahippocampal and inferior parietal cortex at encoding and retrieval. These effects were not as strong as those observed in Post-Meno females, and they were in the opposite direction than the associations seen in Post-Meno females. Taken together with the retrieval effects observed in LV1, our results indicate that better spatial context memory performance at the Pre-Meno stage was observed in females that exhibited increased lateral PFC, angular gyrus and midline cortical activity at encoding and retrieval, and decreased parahippocampal, dorsal occipitotemporal and inferior parietal cortex activity.

In conclusion, we found that both menopause status and chronological age affect spatial context memory behaviorally, though the effect of chronological age is most evident at post-menopause. Menopause status directly affected the direction of age- and performance-related correlations with brain activity in inferior parietal, parahippocampal and occipitotemporal cortices across encoding and retrieval. Moreover, we found that only Pre-Meno females exhibited cortical reinstatement of encoding-related activity in midline cortical, prefrontal, and angular gyrus, at retrieval. This suggests that spatial context memory abilities may rely on distinct brain systems at Pre-compared to Post-Menopause. Future studies should focus on exploring potential mechanisms by which menopause affects age-related differences in memory and brain function. For example, menopause is characterised by a marked decline in centrally circulating estradiol-17β (E2;(Foster, 2012; Harlow et al., 2012; He et al., 2021). Reduced central E2 in ovariectomized rats has been associated with deficits on spatial memory tasks, synaptic loss in the hippocampus and increased neuro-inflammation (Au et al., 2016). Thus, future work exploring the links between blood-based measures of peripheral inflammation, hippocampal volumes, and memory-related brain function may advance our understanding the underlying mechanisms contributing to menopause and age effects on memory-related brain function.

## Acknowledgements

Thank you to the research participants of the Brain Health at Midlife and Menopause (BHAMM) Study for their time and contribution to science. We acknowledge the support of part-time research assistants (H. Azizi, R. Young, A. Condescu, L.Khyatt) and trainees (S. Subramaniapillai, G. Velez Largo, J. Kearley, A. Duval, J. Snytte, S. Loparco) who assisted in participant recruitment or testing or MRI quality control. We are grateful for the support of Team 9, 10 and WSGD Theme of the Canadian Consortium on Neurodegeneration in Aging (CCNA), the MRI Staff at the CIC, DMHUI, and Dr. D. Cohen for help with recruitment. This work was supported by Canada Research Chairs Program (CRC-2022-00240), CIHR Sex & Gender Research Chair (GS9-171369), and NSERC Discovery Grant (RGPIN-2018-05761) awarded to M.N. Rajah.

## Notes

### Competing Interest Statement

The authors have declared no competing interest.

### Summary of Updates

The abstract, significance statement, and introduction was edited. Added new behavioral analysis to examine within group differences in performance by age. Along with minor edits to Tables, Results, and Discussion.

